# Is there a BMAA transfer in the pelagic and benthic food webs in the Baltic Sea?

**DOI:** 10.1101/430330

**Authors:** Nadezda Zguna, Agnes Karlson, Leopold L. Ilag, Andrius Garbaras, Elena Gorokhova

## Abstract

The evidence regarding BMAA occurrence in the Baltic Sea is contradictory, with benthic sources appearing to be more important than pelagic ones. The latter is counterintuitive considering that pelagic primary producers, such as diatoms, dinoflagellates, and cyanobacteria, are the only plausible source of this compound in the food webs. To elucidate BMAA distribution in trophic pathways, we analyzed BMAA in the pelagic and benthic food webs sampled in summer 2010 in the Northern Baltic Proper. As potential BMAA sources, phytoplankton communities in early and late summer were used. As pelagic consumers, zooplankton, mysids and zooplanktivorous fish (herring) were used, whereas benthic invertebrates (amphipods, priapulids, polychaetes, and clams) and benthivorous fish (perch and flounder) represented the benthic food chain. To establish the trophic structure of the system, the stable isotope (δ^13^C and δ^15^N) composition of its components was determined. Contrary to the reported ubiquitous occurrence of BMAA in the Baltic food webs, only phytoplankton and lower consumers (zooplankton and mysids) of the pelagic food chain tested positive. Given that our analytical approaches were adequate, we conclude that no measurable levels of this compound occurred in the benthic invertebrates and any of the tested fish species in the study area. These findings indicate that widely assumed presence and transfer of BMAA to the top consumers in the food webs of the Baltic Sea and, possibly, other systems remain an open question. More controlled experiments and field observations are needed to understand the transfer and possible transformation of BMAA in the food web under various environmental settings.

## Introduction

In the late 1960s, the increased incidence of amyotrophic lateral sclerosis–parkinsonism-dementia complex (ALS–PDC) among native Chamorro population (Guam, Micronesia) was linked to β-N-methylamino-L-alanine (BMAA), a naturally produced non-proteinaceous amino acid (Cox et al., 2003). A causative association between dietary exposure to BMAA and this pathological condition has been broadly discussed, which stimulated research on BMAA production and biomagnification in food webs and development of analytical approaches for detection and quantification of BMAA and its natural isomers, 2,4-diamino butyric acid (DAB), β-amino-N-methyl-alanine (BAMA) and N-(2-aminoethyl) glycine (AEG) (Faassen, 2014; Lance et al., 2018).

The current view is that BMAA supposedly originates from microalgae and cyanobacteria. At first, BMAA production was linked to aquatic and terrestrial cyanobacteria, both free-living and symbiotic, which were considered the only producers of this compound (Cox et al., 2005; Faassen, 2014). BMAA has been detected in a variety of aquatic environments where cyanobacteria blooms can occur, such as oceans, lakes and desert springs, but also in terrestrial environments, like desert mats (Cox et al., 2005, 2009; Craighead et al., 2009; Brand et al., 2010; Li et al., 2010; Metcalf et al., 2015). Recently, several species of diatoms and dinoflagellates were, however, found to also produce BMAA (Jiang et al., 2014a; Jiang and Ilag, 2014). As diatoms and dinoflagellates are the major contributors to primary production in oceans, with their spring and fall blooms being a regular feature in most temperate systems (Klais et al., 2011), the list of potential BMAA sources in aquatic food webs has been greatly expanded as well as the production capacity. These findings have also increased the possible variety of BMAA routes and bioaccumulation pathways in ecosystems (Faassen, 2014).

In various freshwater and marine environments, invertebrate grazers and fish exposed to blooms of potential BMAA producers have also been analyzed and, at least in some studies, found to accumulate BMAA (Brand et al., 2010; Jiao et al., 2014; Andrýs et al., 2015; Réveillon et al., 2015; Salomonsson et al., 2015; Réveillon et al., 2016). However, negative outcomes of such surveys are also quite common (Scott et al., 2009; Niedzwiadek et al., 2012). Then again, most studies reporting BMAA transfer from primary producers to fish have weak sampling design, suffering inconsistencies in both temporal and spatial correspondence between the collected samples of sources and consumers. These inconsistencies hamper quantitative analysis of the BMAA transfer in the food web and make it difficult to trace this compound to specific producers. As a result, controversy surrounds the sources and pathways of BMAA production and accumulation in aquatic ecosystems, which is further complicated by inadequate analytical approaches that have been frequently used in the past.

The evaluation and recent development of the analytical techniques strongly suggest that some of the previous data may overestimate the BMAA concentrations in the environmental samples (Faassen et al., 2012). Indeed, owing to the differences between the analytical methods (Faassen et al., 2012), the reported concentrations even within the cyanobacteria, the most studied group of BMAA producers, vary by orders of magnitude among the studies (Faassen, 2014). Great analytical efforts have been taken to improve and standardize sample preparation and analytical procedures, including using internal standards for accurate identification and quantification of BMAA and isomers in biological samples (Jiang et al., 2013). Currently, the method of choice employs AccQ•Tag Reagent Kit (AQC) derivatization and LC-MS/MS detection in selected reaction monitoring (SRM) mode (Spácil et al., 2010). This configuration has been shown very sensitive and most suitable for analyses of complex samples (Jiang et al., 2014b).

The seasonal cyanobacterial blooms (Bianchi et al., 2000) and detection of BMAA in the Baltic Sea biota raised interest in the BMAA production and fate in this system (Jonasson et al., 2010; Jiang et al., 2014a, 2014b). Current reports of BMAA occurrence in the Baltic Sea region suggest that benthic fish have higher BMAA levels than pelagic ones (Lage et al., 2015) even though no BMAA was found in sediments (Jiang et al., 2014a). Unfortunately, most of the evidence is based on the samples collected outside the spring (diatoms and dinoflagellates) and summer (cyanobacteria) blooms, and in the areas not subjected to intense cyanobacterial blooms, such as the west coast of Sweden (Jiang et al., 2014a; Salomonsson et al., 2015).

One would expect that pelagic fish that feed on zooplankton that is grazing on fresh diatoms, dinoflagellates, and cyanobacteria (i.e., known BMAA producers) in the water column would have higher BMAA concentrations in their body tissues compared to the bottom-dwelling fish that feed primarily on deposit-feeding benthic animals that consume at least partially degraded material. Indeed, even though spring diatom bloom is settled as a relatively fresh material (Bianchi et al., 2002), the microbial activity in the sediment would decrease BMAA content of the algae, and thus generally lower BMAA levels in the benthic food webs. The settling of the summer bloom of cyanobacteria to the sediment has earlier been considered negligible (Sellner et al., 1996), although more recent studies show that other cyanotoxins can accumulate in sediments become transferred to benthic food chains (Mazur-Marzec et al., 2007). To clarify the relative importance of pelagic and benthic pathways of BMAA, data for the relevant sources (i.e., comprising a common food chain), at the proper time period (before and after bloom) or within an ecologically meaningful time period, and from a relatively narrow geographic location for the food chain components in question are needed.

To elucidate BMAA distribution in the Baltic ecological pathways (Fig. 1) before and after a cyanobacteria bloom, we analyzed BMAA levels in the pelagic and benthic food webs using state-of-the-art analytical approach for BMAA detection and quantification. In parallel, to confirm the trophic positions of the food web components used for the BMAA analysis, their stable isotope (δ^13^C and δ^15^N) composition was determined; this is a standard method in ecology and ecotoxicology surveys aiming to elucidate bioaccumulation pathways (Cabana and Rasmussen, 1994).

**Fig. 1.**
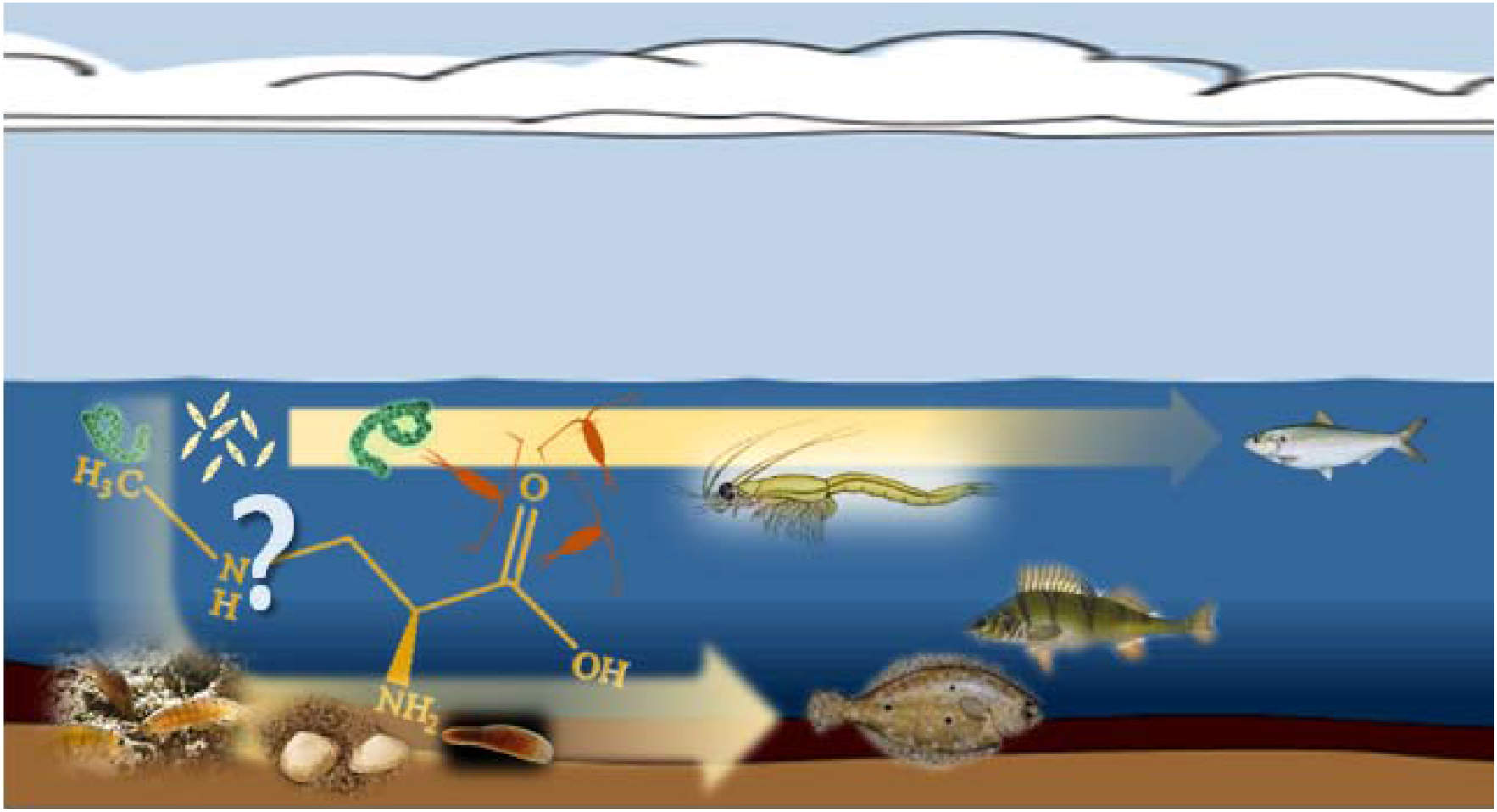
Conceptual diagram of the study design and the potential role of pelagic (invertebrates: zooplankton and mysids, and fish: herring) and benthic (invertebrates: amphipods, clams, priapulids, and polychaetes, and fish: flatfish and perch) food chains in the BMAA transfer from primary producers.

## Material and methods

### Test organisms

As potential BMAA sources, early- and late-summer phytoplankton communities were used. To represent the pelagic chain of consumers, we used mesozooplankton (copepods and cladocerans) and zooplanktivores, mysids (omnivores, feeding on phyto- and zooplankton) and herring (feeding mainly on zooplankton). The benthic chain consumers were invertebrates (amphipods, priapulids, polychaetes, and clams) and nekto-benthivorous fish (perch and flounder) (Table 1; Fig. 1). The same sources and consumers were used for stable isotope (δ^13^C and δ^15^N) composition. All sampled food web components were selected to capture relatively narrow geographic (Fig. 2) and temporal (June-September 2010) variability.

**Fig. 2.**
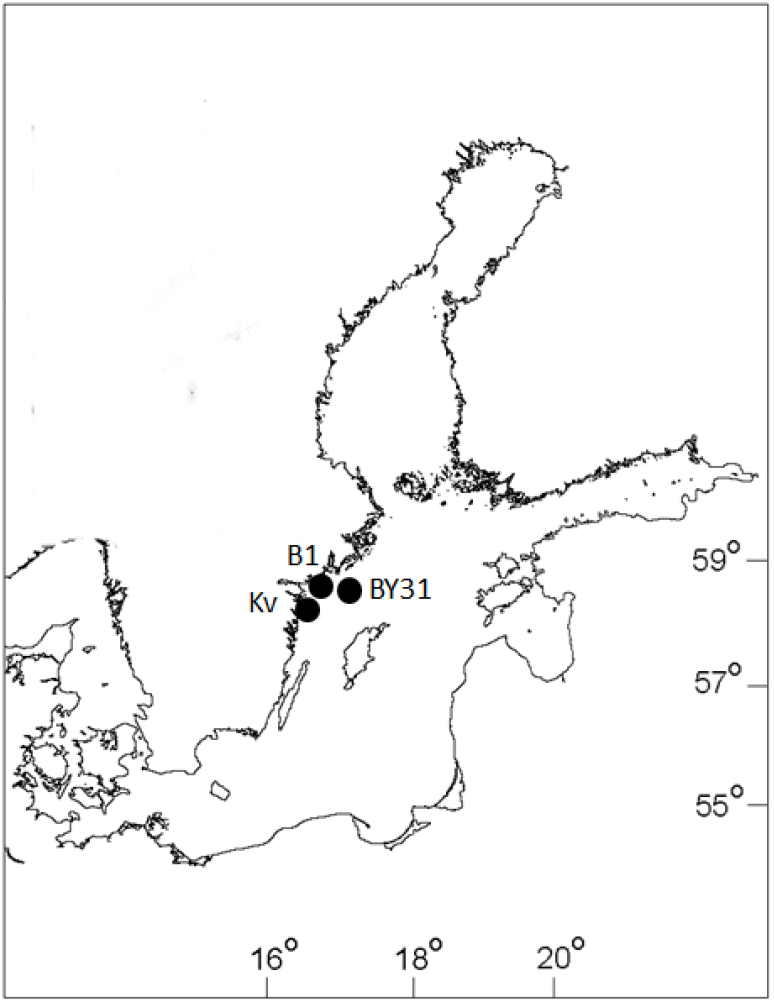
Sampling locations in the norther Baltic Proper for food chain components representing pelagia (BY31, Landsort Deep) and benthic system (B1 and Kvädöfjärden, Kv, both in the viscinity of the Askö Field Station).

Sediment and invertebrates comprising the benthic food chain were collected in a coastal area, and those for the pelagic food chain were collected in the open sea area of the Baltic proper (Fig. 2). The benthic chain samples were collected at several sites in the vicinity of the Askö Field Station (station B1, 58°48’28 N, 17°37’60 E; Swedish National Marine Monitoring Program, SNMMP) in the north-western Baltic proper. Sediment and deposit-feeding macrofauna were collected at stn Håldämman (30 m depth, 58°49’18 N, 17°34’58 E) and stn Uttervik (20 m, 58°50’58 N, 17°32’77 E) on 2–3 sampling events in June and September. We used a benthic sled, set to collect the top 1–2 cm sediment, which was then sieved through a 1-mm sieve to retain macrofauna. The species selected were the bivalve *Limnecola balthica*, the non-indigenous polychaete *Marenzelleria arctia*, the amphipod *Monoporeia affinis*, and the priapulid *Halicryptus spinulosus*. The sieved sediment samples and the recovered animals were frozen and stored at –20 °C until analyses (Table 1).

The pelagic samples (phytoplankton, zooplankton, and mysids) were collected in the Landsort Deep (open-sea stn BY31; 459 m, 58°35’00 N, 18°14’00 E; SNMMP) on 3–4 sampling occasions in June and August-September 2010. Phytoplankton samples were taken using a plastic hose lowered to 20 m depth, giving an integrated 0–20 m sample. After mixing in a bucket, the phytoplankton was filtered on a GF/F (0.7-µm) to gather sufficient material for the analysis. The filters were folded, frozen and stored at –20 °C until analyses. Complementary data on phytoplankton (cell size ≥ 3µm; Figs 3 and 4) taxonomic composition and carbon biomass were obtained from the SHARK database at the Swedish Meteorological and Hydrological Institute (SMHI; *www.smhi.se*). The samples used for the analysis of phytoplankton community structure were collected within 3–4 days of our sampling occasions, and the community composition was assumed to represent that in our samples analyzed for BMAA. Zooplankton were collected by vertical tows in the upper 30 m using a 90-µm WP-2 net (diameter 57 cm). Species that dominated, the copepods (*Acartia* spp. and *Eurytemora affinis*) and the cladocerans (*Bosmina coregoni maritima*), were picked under a stereomicroscope to avoid contamination with phytoplankton and frozen in bulk. The rest of the single-tow sample was preserved in 4% borax buffered formaldehyde for species identification and community analysis. Mysids (*Mysis mixta* and *N. integer*) were collected in the upper 100 m using either a 500- or a 200-µm WP-2 net (diameter 57 cm), length-measured, and frozen individually in Eppendorf tubes (Table 1).

**Fig. 3.**
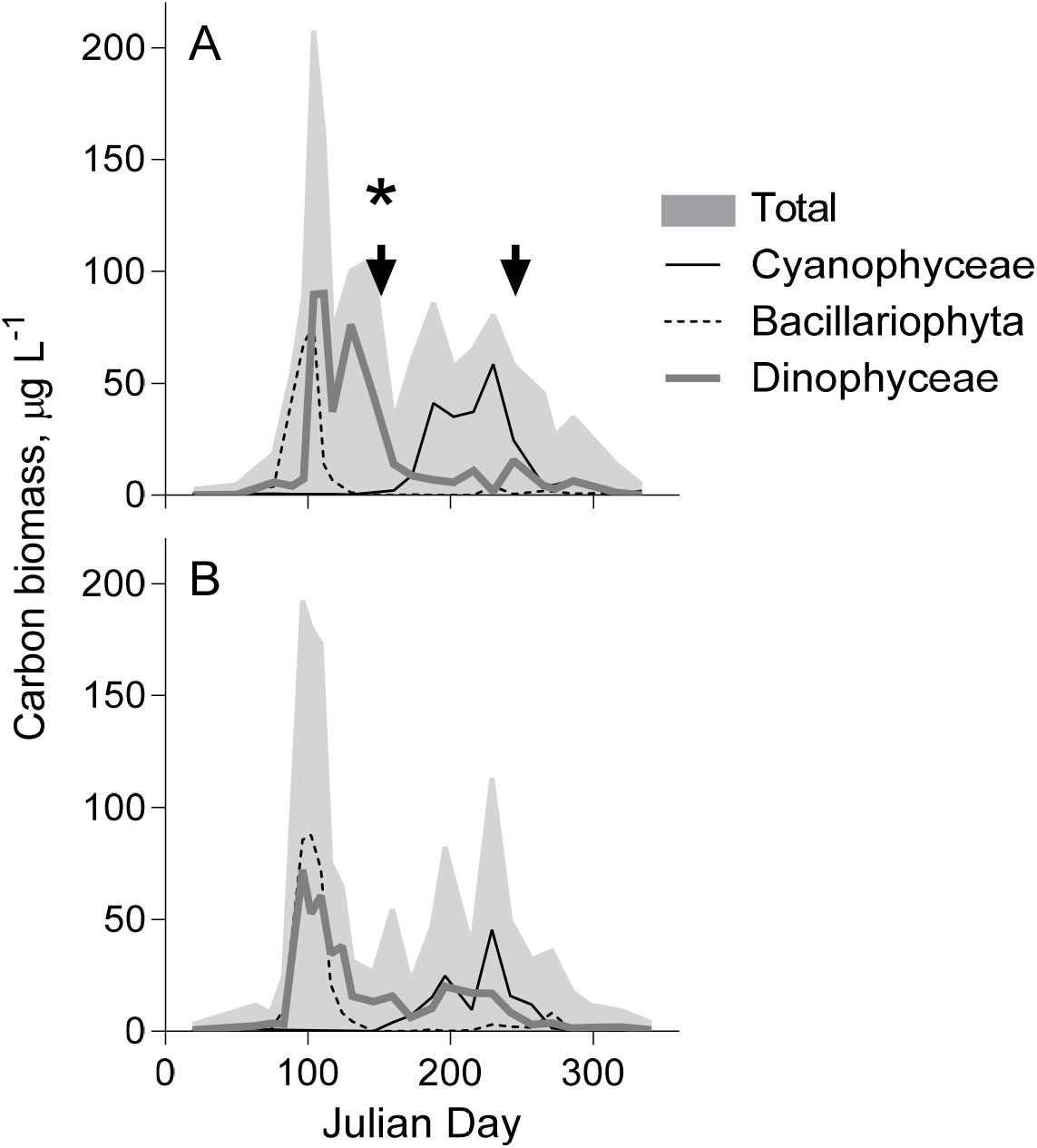
Phytoplankton dynamics in the study areas: (A) the Landsort Deep (pelagic food chain) and (B) Askö station (benthic food chain). The data on the total carbon biomass of phytoplankton and seasonal dynamics of the main BMAA producers (cyanobacteria, diatoms and dinoflagellates) in the year 2010 are obtained from the SHARK database. Sampling occasions are indicated by arrows and the BMAA-positive sample with asterisk.

**Fig. 4.**
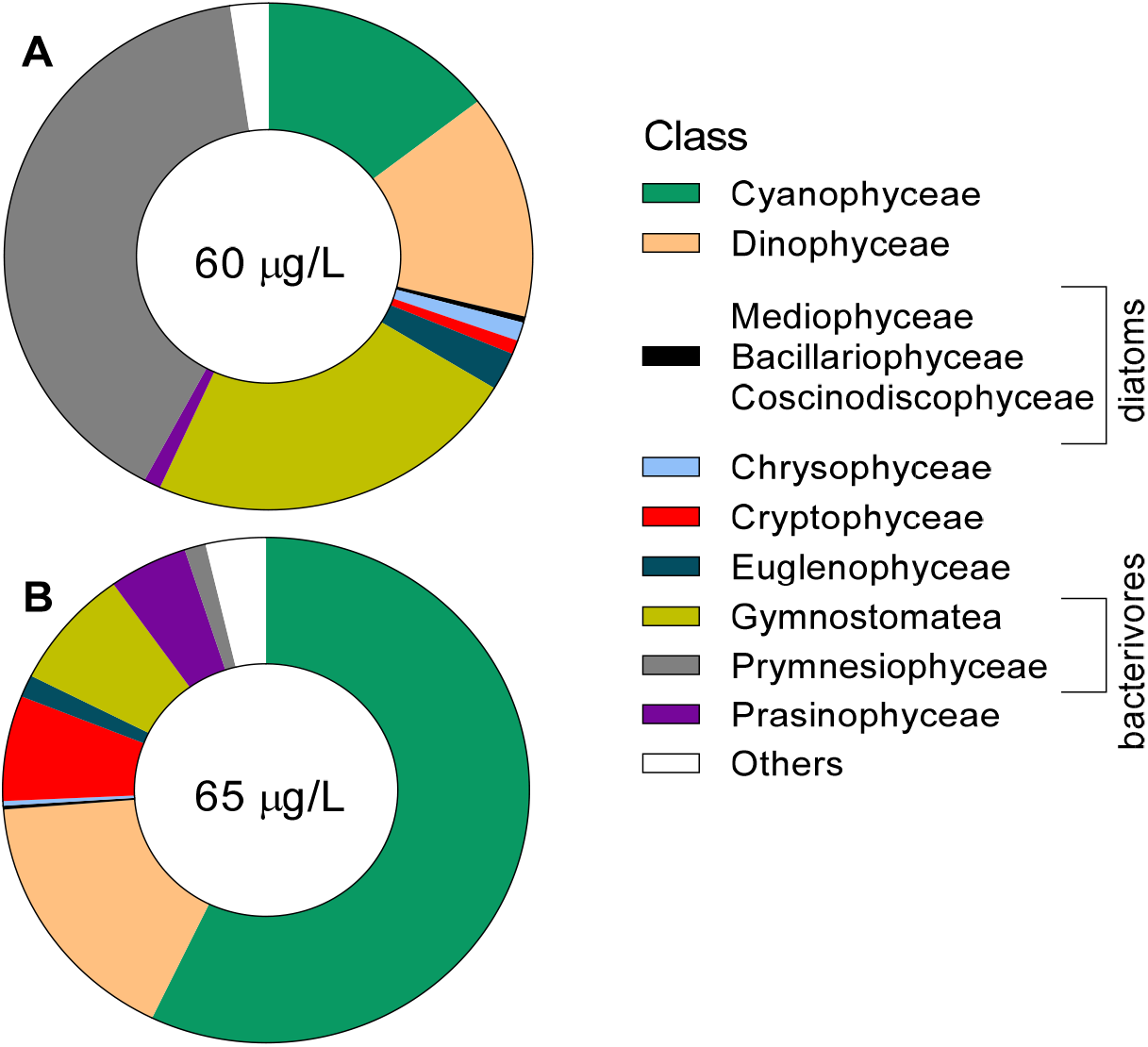
Composition of the phytoplankton communities that were (A) BMAA-positive (June) and (B) BMAA-negative (August). Number in the middle of the pie diagram is the total carbon biomass of the phytoplankton at the time of collection.

Fish were obtained from the Swedish Museum of Natural History; the collection was made in September 2010 as a part of the Swedish National Monitoring Program for Contaminants in Marine Biota. Bottom-feeding fish species included the European flounder *Platichthys flesus* (Aarnio et al., 1996) and the Eurasian perch *Perca fluviatilis* (Mustamäki et al., 2014); these were sampled in Kvädöfjärden (58.0497° N, 16.7831° E). As a fish with pelagic feeding, we used the Baltic herring *Clupea harengus* (Casini et al., 2004) sampled in the Landsort Deep. All fish was length-measured, the epidermis and subcutaneous fatty tissue are carefully removed, and muscle tissues were taken from the middle dorsal muscle layer for BMAA and stable isotope measurements (Table 1).

### Positive controls

Two positive controls were used: (1) a blue mussel (*Mytilus edulis*; Bivalvia) sample that tested positively in the previous study (Jiang et al., 2014b) and (2) cyanobacterium *Nodularia spumigena* (strain AV1) grown axenically in culture and harvested at exponential and stationary growth phases. The cyanobacterium was grown in a modified Z8 nutrient solution in 16:8 h light:dark regime (40 µE cm^−2^ s^−1^) at 20–21 °C with constant shaking and harvested when cell density reached approximately 5 × 10^6^ cell/mL (3–5 days) and 10^9^ cell/mL (9–10 days) for the exponential and stationary phase samples, respectively; three replicates for each group were obtained.

### BMAA analysis

Samples for the UPCL-MS/MS analysis of BMAA were prepared according to Jiang and co-workers (Jiang et al., 2013; 2014), with some modifications. The sample material was homogenized with a mortar and pestle with the addition of liquid nitrogen. A weighed subsample of the homogenate (20 to 50 mg; Mettler Toledo XP6; +/− 0.001 mg) was mixed with 600 µL of water and subjected to ultrasound treatment with Vibra CellTM, Sonics & Materials Inc. Danbury CT, USA (3 min, 1 second on/off pause, 70% amplification) in an ice-water bath. An aliquot corresponding to 10-mg of the homogenate was transferred to a glass hydrolysis vial together with 10 µL of deuterated BMAA standard, water, and 6M HCl, and hydrolyzed at 110°C for 20 h.

The hydrolyzed samples were filtered in a centrifuge using spin filters (0.2 µm, Thermo Scientific, USA) and subjected to a two-step clean-up: (1) liquid-liquid extraction with chloroform to remove lipids and other hydrophobic compounds, and (2) solid phase extraction (SPE) with Isolute HCX-3 column (Biotage Sweden AB) using 1 mL of 0.1% formic acid in water and 1 mL of methanol for column conditioning followed by sample application and washing by 1 mL of methanol and 1 mL of 0.1% formic acid in water. For elution (two rounds), 0.8 mL of 2% NH4OH in methanol were used.

Samples were derivatized with 6-aminoquinolyl-Nhydroxysuccinimidyl carbamate (AQC) reagent using AccQ·Tag kit (Waters; WAT052880, Milford, USA), reconstituted in 20 µL 20 mM HCl, and mixed with 60 µL borate buffer (from AccQ·Tag kit) and 60 µL of AQC reagent. Before analysis, all samples were dried and reconstituted in the initial UHPLC mobile phase (30 µL of 5% acetonitrile in water). All samples were analyzed in duplicates.

The samples were analyzed by UHPLC-MS/MS as described elsewhere (Jiang et al. 2012) using TSQ Vantage triple quadrupole mass spectrometer (Thermo Fisher Scientific, USA) equipped with an Accela pump and auto-sampler as well as a degasser. An additional pump (Rheos 4000 pump, Flux instruments) was used for post-column addition of 0.3% acetic acid in acetonitrile at the flow rate of 600 µL/min. The UHPLC system was equipped with an ACCQ-TAG™ ULTRA C18 column (100×3×2.1 mm, 1.7 mm particle size, Waters, Ireland). Binary mobile phase (MP) was delivered according to the programmed gradient (MP A: 0.3% acetic acid and 5% acetonitrile in water and MP B: 0.3% acetic acid in acetonitrile) as follows: 0.0 min, 0% B (flow rate 200 µL/min); 10.0 min, 10% B (flow rate 400 µL/min for the rest of the program); 11.0 min, 80% B; 12.0 min, 80% B; 12.1 min, 0% B; and 16.0 min, 0% B. MS/MS analysis was performed by multiple reaction monitoring (MRM) in positive ion mode. One monitored MRM transition (459.18 > 119.08) was general for BMAA and its isomers (BAMA, DAB & AEG), whereas other three MRM transitions were diagnostic for the particular isomers (459.18 > 258.09 for BMAA and BAMA, 459.18 > 188.08 for DAB, and 459.18 > 214.10 for AEG), and one MRM transition was unique for d3-BMAA internal standard (462.20 > 122.10). The unique transitions in combination with carefully compared retention times and peak area ratio between the general and diagnostic transitions allowed a reliable detection and quantification of BMAA.

Method performance was evaluated using spiked samples prepared with dried and pulverized cod muscle (negative control); the latter contained no BMAA as shown in previous studies (Faassen et al., 2012; Jiang et al., 2014b). The lowest concentration spiked was 0.05 ng/10 mg tissue wet weight, and this amount was detected S/N>10. Also, a calibration curve for the range of concentrations between 0.1–10 ng BMAA on column showed excellent linearity. Limit of detection (LOD) and limit of quantification (LOQ) were estimated to be below 0.01 mg BMAA/g wet weight by analyzing spiked samples of crayfish muscle tissue. The accuracy and precision of the method were evaluated with the quality control samples at different BMAA concentrations and were determined to be 108–119% and <15% RSD, respectively. The sensitivity of the method was determined as low as 4.2 fmol BMAA/injection (Jiang et al., 2014b). Additionally, we conducted a recovery test by spiking the matrix of cod muscle; this material was proven to be negative for BMAA in the previous study (Jiang et al., 2014). The spiked sample was analyzed in triplicate, and the recovery was 106%, which is in agreement with the previously published method performance report (Jiang et al., 2014b).

### SIA analysis

All biological material and sediment were dried to a constant weight (24 to 60 h, depending on the sample type) at 60 °C. To prepare the samples of zooplankton, phytoplankton, and sediment, the bulk material was used, whereas, for the samples of fish and invertebrates, we subsampled ∼0.7 mg dry weight (DW). The samples were transferred to tin capsules, dried again at 60 °C for 24 h, and shipped to the Center for Physical Science and Technology, Vilnius, Lithuania. The SIA was conducted using Flash EA 1112 Series Elemental Analyzer connected via a Conflo III to a DeltaV Advantage isotope ratio mass spectrometer (all Thermo Finnigan, Bremen, Germany). Ratios of^14^N:^15^N and^12^C:^13^C are expressed relative to the international standards, atmospheric air (N) and Pee Dee Belemnite (C) and presented in a δ-notation, parts per thousand difference from the standard. Internal reference (cod muscle tissue) was analyzed every 20 samples; the analytical precision was 0.1 ‰ for both δ^15^N and δ13C.

## Results and Discussion

Contrary to the reported ubiquitous occurrence of BMAA in the Baltic food webs, only phytoplankton (seston), zooplankton and mysid samples tested positive. None of the benthic invertebrates or fish species were found to contain BMAA. Moreover, no measurable BMAA levels were detected in the sediment. Given that the analytical performance was adequate, and both positive controls, i.e., the blue mussel and the cyanobacterium, tested positive, we conclude that all samples that tested negative contained no measurable levels of this compound.

The blue mussel positive control yielded 3.1 µg g^−1^ wet weight, which is similar to the previously obtained BMAA concentration in that sample (Jiang et al., 2014b). In the *Nodularia spumigena* harvested at the exponential and stationary phase, the BMAA concentration (mean ± SD, n = 3) was 0.98 ± 0.06 and 0.13 ± 0.02 µg g^−1^ wet weight, respectively. These concentrations are comparable to those reported for this and other *Nodularia* species (Cox et al., 2005). Thus, both types of the control material were found to contain the expected amounts of BMAA. Moreover, in the cyanobacterium, the BMAA levels were approximately 8-fold higher during the exponential growth compared to the aging culture.

The BMAA concentrations measured in seston and zooplankton varied from 0.83 to 1.13 µg g-1 wet weight, which corresponds to approximately 7.5–9.5 µg g^−1^ dry weight using common conversion factors for phyto- and zooplankton (Kiørboe, 2013; Sládeček and Sládečková, 1963). In the Baltic Sea, the BMAA concentration in cyanobacteria-rich phytoplankton community has been reported to vary from 0.001 to 0.015 μg BMAA/g dry weight, and that of zooplankton 0.004 to 0.087 μg BMAA/g dry weight (Jonasson et al., 2010). These values are more than 100-fold lower than the concentrations measured in our study, but also several orders of magnitude lower than the reported BMAA concentrations in plankton collected in other systems. For instance, samples of cyanobacteria in Dutch urban waters (Faassen et al., 2009), British (Metcalf et al., 2008) and South African (Esterhuizen and Downing, 2008) lakes as well as Taihu lake in China (Jiao et al., 2014) are in the µg BMAA/g dry mass range or higher, which is two to five orders of magnitude higher than the range reported by Jonasson and co-workers (Jonasson et al., 2010). A reason for these conflicting results can be related to the fact that BMAA quantification in the latter study was based on a semi-quantitative method, with no internal standard; moreover, no LOD values were reported. The method that was employed involves a clean-up procedure based on the liquid-liquid extraction followed by a solid phase extraction, which can introduce a significant loss of BMAA during the work-up. As shown by the inter-laboratory evaluation, the recovery of this clean-up procedure might be as low as <10% (Faassen et al., 2016). Therefore, the addition of an internal standard before the clean-up is required to control for the losses during the sample work-up.

Only a single phytoplankton sample was found to contain BMAA, and the concentration measured was within the BMAA levels reported in studies that are considered as truly reliable with regard to the quality assurance and the protocols involved (Lance et al., 2018). The phytoplankton samples were taken during two periods: (1) past the spring bloom comprised by diatoms and then dinoflagellates but before the onset of the summer cyanobacteria bloom (June), and (2) after the cyanobacteria bloom collapse (August; Fig. 3A). Both spring and summer blooms were of average magnitude and duration in relation to the multiyear variability (Klais et al., 2011; Wasmund, 1997) and similar between the coastal and open sea areas (Fig. 3). However, only one phytoplankton sample taken in the mid-June contained BMAA (Fig. 4A, Fig. 5). The phytoplankton community composition at the time of sampling was typical for this area and the time of year, with biomass composed of ∼40% of haptophytes, 23% of ciliates, 16% cyanobacteria, 15% dinoflagellates, the rest being a mixture of summer species (Fig. 4A). Thus, there was a high contribution of bacterivores (ciliates and haptophytes, 64% together) to the community. The August samples that did not yield measurable BMAA quantities had approximately 3.5-fold higher percentage of cyanobacteria (Fig. 4B) and 8-fold lower percentage of bacterivorous species. In no case contribution of diatoms exceeded 0.5%. Given this variability in the taxonomic composition of the phytoplankton, with two potentially significant BMAA producers (i.e., cyanobacteria, and dinoflagellates) being moderately abundant, we cannot identify a plausible source of the BMAA in our sample. It is also possible that during the decline of the bloom, the cyanobacteria have lower BMAA levels as what we found in our positive control samples collected during the stationary phase; this might have contributed to the non-detectable levels in the August sample.

**Fig. 5.**
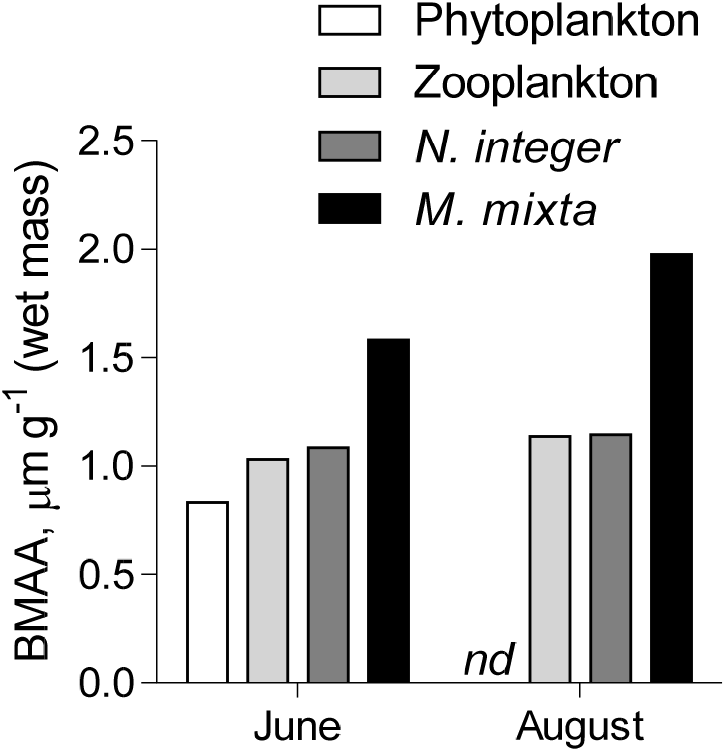
BMAA concentration in phytoplankton, zooplankton, and mysids (*N. integer* and *M. mixta*) collected before (June) and after (August) the cyanobacterial bloom. All samples were composite to integrate respective food web components over several sampling occasions; *nd* – not detectable.

It is also possible that heterotrophic protists can accumulate BMAA and contribute to its bulk concentration in seston. It has been suggested that picocyanobacteria, such as *Synechococcus*, can contribute substantially to BMAA production and transfer to primary consumers (Masseret et al., 2013). In the Baltic microbial loop, the bacterivorous taxa (ciliates and mixotrophic phytoplankton) feeding on picocyanobacteria would then have a high capacity to accumulate BMAA and transfer it further to the primary consumers, such as zooplankton (Motwani and Gorokhova, 2013). Our findings support this pathway, because the high contribution of bacterivores coincided with the presence of BMAA in the seston sample, whereas neither absolute nor relative amount of filamentous cyanobacteria contributed to the BMAA levels (Fig. 4A). Analyzing more samples with different community structure and linking BMAA levels to the community composition in the microbial loop would be one way to identify the main BMAA producers and consumers in these communities. Another approach would be to use culture-based studies with species that were putatively identified as the BMAA producers in the field.

Zooplankton and mysid samples were found to contain BMAA, both before and after the cyanobacterial bloom, despite the lack of BMAA in the phytoplankton sampled in August (Fig. 5). The composition of zooplankton communities differed substantially between the sampling occasions, with cladocerans comprising > 60% of the total zooplankton biomass in June and copepods dominating (>70%) in August (data not shown). Moreover, BMAA levels in mysids differed between the species, being 50–70% higher in *M. mixta* than in *N. integer*. The difference could be related to the more herbivorous diet of *N. integer* and more zooplanktivorous feeding of *M. mixta* (Rudstam et al., 1989). In line with this, the δ^15^N values were 1.5‰ lower in *N. integer* compared to *M. mixta* (Fig. 6A), indicating more herbivorous diet. Therefore, the stepwise increase in BMAA from phytoplankton to *M. mixta* (Fig. 5) could be indicative of biomagnification, albeit more data are needed to substantiate this suggestion.

**Fig. 6.**
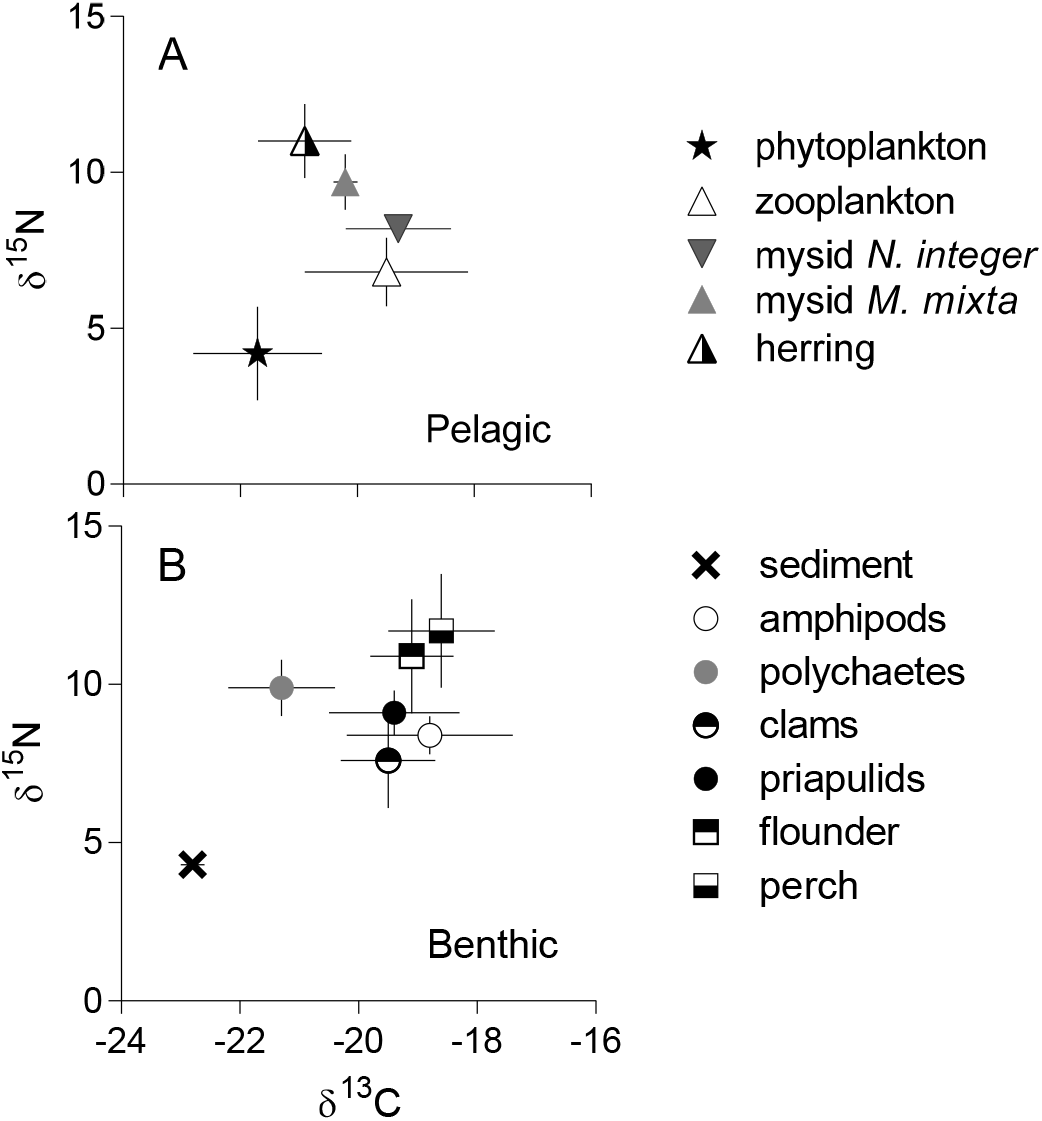
Food web structure determined by SIA (δ^15^N vs. δ^13^C; mean ± SD); pelagic (A) and benthic (B) compartments. See Table 1 for details on the food web components, sample origin, and the number of samples analyzed.

It has been repeatedly stressed that when collecting material for bioaccumulation studies, it is essential to use an adequate temporal and spatial resolution and to ensure the pathway identification and trophic relatedness of the ecosystem components (Walters et al., 2016). To confirm the trophic relationships between the organisms involved in the bioaccumulation assessment, a stable isotope approach has been commonly used (Cabana and Rasmussen, 1994). We applied SIA to confirm the trophic linkages for the consumers, particularly the fish species with a non-sedentary behavior. The SIA data (Fig. 6) confirmed that all animals analyzed for BMAA were occupying the trophic positions as expected (based on their δ^15^N values) and belonged to the same food web (δ^13^C values). In particular, the isotopic signatures supported the assumption that herring collected in the area where zooplankton and mysids were collected was indeed relying on these prey (Fig. 2); however, we found no evidence of the BMAA presence in the fish. As no BMAA was found in the sediment and benthic invertebrates (four species), it is not particularly surprising that all benthic fish that were analyzed (two species) were negative.

The current view that in the food webs BMAA may be transferred from producers to zooplankton and other filter-feeders, such as bivalves, and bioaccumulated in fish feeding on these invertebrates is based only on a limited data. In particular, these patterns have been reported using material collected in the Baltic Sea (Jonasson et al., 2010), the Florida Bay, USA (Brand et al., 2010), and the Portuguese transitional waters (Lage et al., 2014). Of these studies, only the latter used a controlled sampling design, with the consumers and primary producers collected systematically from the same environment. The authors have found a correlation between dinoflagellate *Gymnodinium catenatum* blooms and a subsequent, with a few-day lag phase, an increase of BMAA concentration in the river cockles. They have also used a laboratory experiment to demonstrate BMAA production by *G. catenatum*. In other studies, the sample material originated from various time periods and sampling locations with no clear trophic linkage between the producers and consumers. This holds true for the samples of plankton, fish and benthic invertebrates analyzed by Jonasson and co-workers (Jonasson et al., 2010), which originated from the geographically distant regions and food webs, i.e., the Baltic proper and the North Sea, which makes it questionable for the bioaccumulation assessment.

Although the bioaccumulation/biomagnification of BMAA has been broadly assumed, the *in situ* evidence using material collected in a systematic way is still limited. Our findings indicate that ubiquitous transfer of BMAA to the top consumers in the food webs of the Baltic Sea and, possibly, other systems, is questionable. It also implies that invertebrates and fish associated with benthic food sources in this system are less likely to accumulate BMAA compared to the pelagic food chain. However, some tissue effects on BMAA accumulation may occur resulting in higher values in, for example, protein-rich tissues/organisms. In line with this, results from the most reliable studies show that marine bivalves are to date the matrix containing the highest amount of BMAA, far more than most fish muscles, but with an exception for shark cartilage (Lance et al., 2018). Therefore, to understand BMAA bioaccumulation, the concentrations on the protein basis might need to be used.

Our findings suggest that BMAA levels in the Baltic food webs are much below the suggested risk level considering effect concentrations reported for invertebrates (Esterhuizen-Londt et al., 2015; Faassen et al., 2015; Lepoutre et al., 2018). Moreover, all analyzed fish species, both zooplanktivorous and benthivorous, tested negative, which implies a low risk for the top consumers, such as seal, birds, and humans. However, our sampling was limited to a few occasions in the summer and a relatively small geographic area, whereas it is possible that BMAA levels vary depending on environmental conditions and ecological responses of its producers. Therefore, to understand the magnitude of this variability, it is important to conduct a systematic sampling throughout a year and in different parts of the Baltic Sea. Sensitive and validated analytical methods should be used to ensure that the obtained results are consistent. Quality control samples must be included in the surveys to evaluate the performance of the methods, particularly the analyte recovery and accuracy of the results. Finally, more controlled experiments and field observations are needed to understand the trophic transfer and possible transformation of BMAA in various environmental settings.

---

**Table S1.**
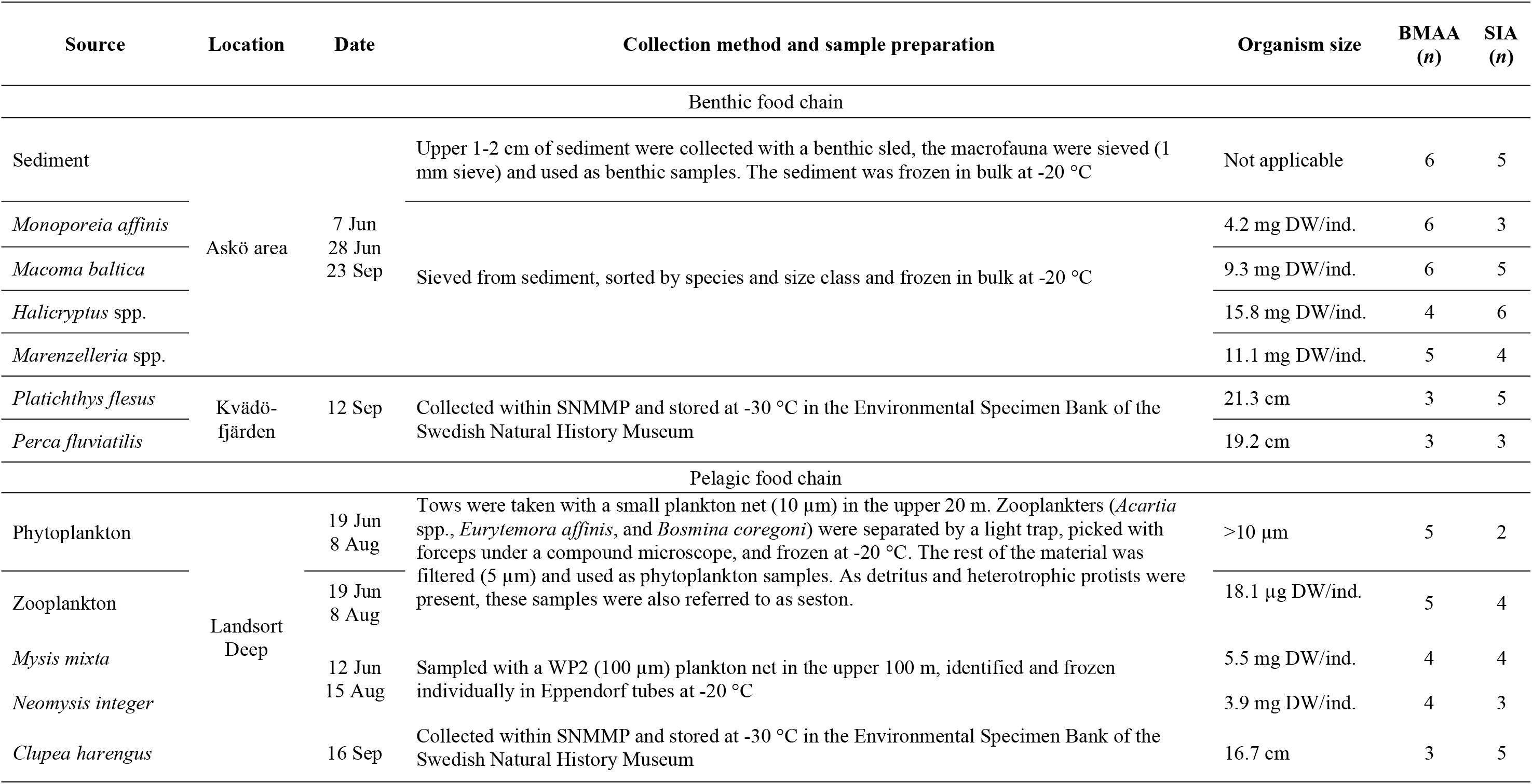
Summary of samples used for BMAA analysis and SIA and collection methods. Organism size for the benthic invertebrates and mysids was determined as the average dry mass per individual in the SIA samples and the number of individuals used to prepare these samples, and for the fish, it is the total body length. For BMAA, on each sampling occasion, 2-3 replicate samples for invertebrates and 3 replicates for fish were analyzed; at least two technical replicates were used.

